# Super-resolution mapping of anisotropic tissue structure with diffusion MRI and deep learning

**DOI:** 10.1101/2023.04.04.535586

**Authors:** David Abramian, Anders Eklund, Evren Özarslan

## Abstract

Diffusion magnetic resonance imaging (diffusion MRI) is widely employed to probe the diffusive motion of water molecules within the tissue. Numerous diseases and processes affecting the central nervous system can be detected and monitored via diffusion MRI thanks to its sensitivity to microstructural alterations in tissue. The latter has prompted interest in quantitative mapping of the microstructural parameters, such as the fiber orientation distribution function (fODF), which is instrumental for noninvasively mapping the underlying axonal fiber tracts in white matter through a procedure known as tractography. However, such applications demand repeated acquisitions of MRI volumes with varied experimental parameters demanding long acquisition times and/or limited spatial resolution. In this work, we present a deep-learning-based approach for increasing the spatial resolution of diffusion MRI data in the form of fODFs obtained through constrained spherical deconvolution. The proposed approach is evaluated on high quality data from the Human Connectome Project, and is shown to generate upsampled results with a greater correspondence to ground truth high-resolution data than can be achieved with ordinary spline interpolation methods.

## Introduction

Diffusion magnetic resonance imaging (diffusion MRI) is an MRI modality used to probe and characterize the diffusion process of water molecules within the tissue. Diffusion MRI has proven sensitivity to many diseases affecting the tissue, making it an indispensable tool in diagnostic medicine.^1, 2^ Diffusion of molecules is influenced by the ambient environment, which imprints its signature on the diffusion MRI signal. Of particular interest to us are the microstructural characteristics of the tissue that influence the diffusion MRI signal, which could be instrumental in not only improving the diagnostic utility of MRI, but also address rather fundamental questions regarding the make up and organization of the tissues.^3^

For example, one of diffusion MRI’s principal uses involves estimating the local orientation of axonal fibers within white matter in the brain. These, in turn, can be used to estimate the long-range connections inside the brain, through a procedure known as *tractography*^4–6^, which consists of alternately sampling the local fiber orientation at a given position in the brain and taking a small step in that direction. Performing this procedure starting from multiple points within the brain makes it possible to reconstruct the pathways of the axonal bundles inside the brain.

There are multiple ways of describing and summarizing the local diffusion properties of water molecules, which differ in the amount of precision they afford, the complexity of the acquisition process required to estimate them, and their suitability for various purposes. Diffusion tensor imaging (DTI)^7^ is the most common such method, in which a tensor is fitted to the diffusion signal at every brain voxel. For voxels containing a single fiber population, this provides a suitable estimation of its primary orientation, but the method is incapable of properly representing diffusion along multiple crossing fiber populations.

To address this challenge, a number of higher order models have been proposed over the years.^8–17^ Among these, the constrained spherical deconvolution (CSD) method^15^ has been widely employed in tractography studies. In this approach, the signal is envisioned to arise from a continuous distribution of fibers. The associated density function is referred to as the “fiber orientation distribution function (fODF),” which is estimated by deconvolving the detected signal with a response function taken to be the same for all fibers. Such models rely on diffusion MRI data with a relatively large number of diffusion-weighted MRI acquisitions each having a measurement sensitized to diffusion taking place along a different orientation.^18^ Moreover, the data are to be acquired preferably at a larger diffusion-weighting, resulting in a reduced signal-to-noise ratio (SNR) compared to what can be achieved in DTI acquisitions. In order to address these issues, spatial resolution is typically reduced. Having voxels of size 2-3 mm is thus common in conventional diffusion MRI acquisitions.

To overcome this serious limitation, we introduce a deep learning based super-resolution technique that provides an accurate upsampling of data collected for CSD analysis, thus providing fODF-valued images with voxels considerably smaller than the resolution employed in data acquisition.

Adaptation of deep learning for achieving super-resolution has been a popular research topic during recent years. Deep learning methods have already been incorporated into clinical acquisitions for generating high-resolution reconstructions of images accounting for space-dependent noise patterns^19^. Most of the work has focused on two-dimensional (2D) images^20–22^, and much less work has extended the methods from 2D to 3D^23–26^. However, diffusion MRI data is 4D, as many volumes (e.g. 7 - 300) are collected with diffusion weighting along different directions and with different b-values. However, existing deep learning frameworks do not support 4D convolutional neural networks (CNNs). Since the order of the collected volumes is not important, compared to functional MRI which has a time dimension, it is sufficient to use a multi-channel 3D CNN. However, doing so would require too much GPU memory to work on the full dataset. To use super resolution techniques for diffusion MRI data can therefore be done in different ways; independently for a small number of adjacent 2D slices, independently for a small number of 3D volumes, or independently for all measurements or coefficients in a small number of adjacent voxels. While the first two approaches can use a larger spatial context, they will run into GPU memory problems when using many measurements or coefficients at the same time.

Qin et al.^27^ used deep learning to obtain super-resolved diffusion MRI data. The input in their framework involved all measurements for a small number of voxels. There are a number of differences compared to our work. First, they apply the super-resolution to raw diffusion measurements, or more precisely a learned sparse dictionary, while we apply it to 45 coefficients in a spherical harmonics (SH) representation of the fODFs obtained through CSD. Second, their network generates high-resolution maps of scalar parameters representing some microstructural features (orientation dispersion, intra-cellular volume fraction and cerebrospinal volume fraction) while we generate a high-resolution version of the 45 SH coefficients. Our work is thereby more aimed at tractography. Luo et al.^28^ instead used a generative adversarial network (GAN) architecture to directly upsample low resolution diffusion MRI data to a higher resolution. Although a 3D CNN was used inside the GAN, each sub-volume contained only 11 × 11 × 11 voxels to be able to fit all 95 measurements at the same time. None of the previous works on this topic performed repeated upsampling.

## Methods

All data processing was done using custom Python (version 3.9) scripts and Jupyter notebooks. All processing of diffusion MRI data was done using the dipy toolbox^29^ (version 1.5) for Python. Deep learning models were implemented in TensorFlow (version 2.4.1).

### Data

Data used for all analyses were obtained from the WU-Minn Human Connectome Project (HCP)^30^. We use the 100 unrelated adult subject sub-group (54% female, mean age = 29.11 years, range = 22–36). Five of the subjects were excluded due to incomplete WM coverage of the diffusion MRI data, leaving a total of 95 subjects. The data have been previously subjected to a minimal preprocessing pipeline^31^. The HCP diffusion MRI data have isotropic voxels of size 1.25 mm, and consist of 288 diffusion-weighted volumes, with 3 shells of data with b-values of 1000, 2000 and 3000 s/mm^2^ and 90 diffusion directions each, in addition to 18 volumes collected with no diffusion weighting. Besides these 18 volumes, only the volumes in the 3000 b-value shell were used, as the fitting was done with a single-shell model, and constrained spherical deconvolution (CSD) benefits from a strong diffusion encoding.

In order to simulate low resolution data, the original diffusion MRI volumes were downsampled by averaging the signal in each non-overlapping 2 × 2 × 2 voxel region. Given that in MRI acquisition the signal within a voxel integrates the contributions from all components within the voxel, such a downsampling approach mimics the effects of having acquired MRI data with half the spatial resolution in each axis.

The HCP data include a parcellation volume for each subject, where each brain voxel is classified as belonging to one of a number of anatomical regions and tissue types. These tissue labels were used in order to evaluate the performance of the upsampling methods on different tissue types, namely white matter (WM), gray matter (GM) and cerebrospinal fluid (CSF).

### Data processing

For every subject, fODFs were estimated at every voxel by fitting a constrained spherical deconvolution (CSD) model^15^. The WM response function used for deconvolution, representing the expected measured diffusion signal for a single fiber bundle, was estimated from highly anisotropic voxels in and near the corpus callosum. The fODFs were transformed into the spherical harmonic (SH) domain, which constitutes a natural representation for spherical functions. The SH basis used was the one employed by Descoteaux et al. in their work on Q-ball imaging^32^, with a maximum order of 8, resulting in 45 real SH coefficients per voxel. The use of the SH domain has several advantages over using raw data or the fODFs spherical functions directly. First, it allows a full description of the spherical functions using a limited number of coefficients. Furthermore, it provides a generic domain independent from the specific sampling scheme chosen for the fODFs, and which can likewise be calculated from any chosen spherical sampling. It is therefore possible, at least in theory, to use our trained super-resolution network on diffusion data collected with another sampling scheme.

Due to the differential proton-density and magnetic relaxation effects over tissue types, fODFs generated in this way do not correspond to true probability distributions, i.e., their integral over the surface of the sphere is not equal to unity. In order to obtain true probability distributions, the SH coefficients at each voxel were normalized by dividing them by 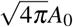, where *A*_0_ represents the 0-th SH coefficient. This has the added advantage of requiring one fewer SH coefficient, as *A*_0_ becomes 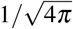 for all voxels.

The described procedure was carried out separately on the low-resolution and high-resolution diffusion MRI data.

### Neural network model

We used a fully-connected neural network model to upsample the low-resolution SH data to its original resolution. The network was provided with SH coefficients from a cubic region of the low-resolution data as input, and was trained to output the SH coefficients of the 8 high-resolution voxels corresponding to the central voxel of the low-resolution input region. Specifically, the number of input dimensions are 1188 (3 x 3 x 3 voxels for 44 SH coefficients, flattened to a vector) and the number of output dimensions are 352 (a vector which can be reshaped to 2 x 2 x 2 voxels for 44 SH coefficients).

The neural networks consisted of several fully-connected layers, with batch normalization and ReLU activation functions for the hidden layers, and no activation function for the final layer. All networks were trained with mean squared error loss, Adam optimizer^33^, and a batch size of 512 for 100 epochs. A grid search was used to select the number of fully-connected layers and the number of nodes per layers. It was found that 3 layers of 1000 nodes each resulted in a good trade-off between validation performance and degree of overfitting. Figure 1 presents a schematic representation of the employed network. Networks were trained on SH data from 50 subjects, with 10 subjects used for validation, and the remaining 35 for testing. Data were split on the subject level, to avoid data leakage^34^. As every low-resolution brain voxel constitutes a datapoint, and with an average of 130000 low-resolution voxels per subject, this resulted in 6.7 million, 1.4 million, and 4.7 million training, validation, and test data points, respectively. The final network has a total of 3.5 million trainable parameters, and the training took approximately 90 minutes on an Nvidia Quadro RTX 8000 GPU.

**Figure 1.**
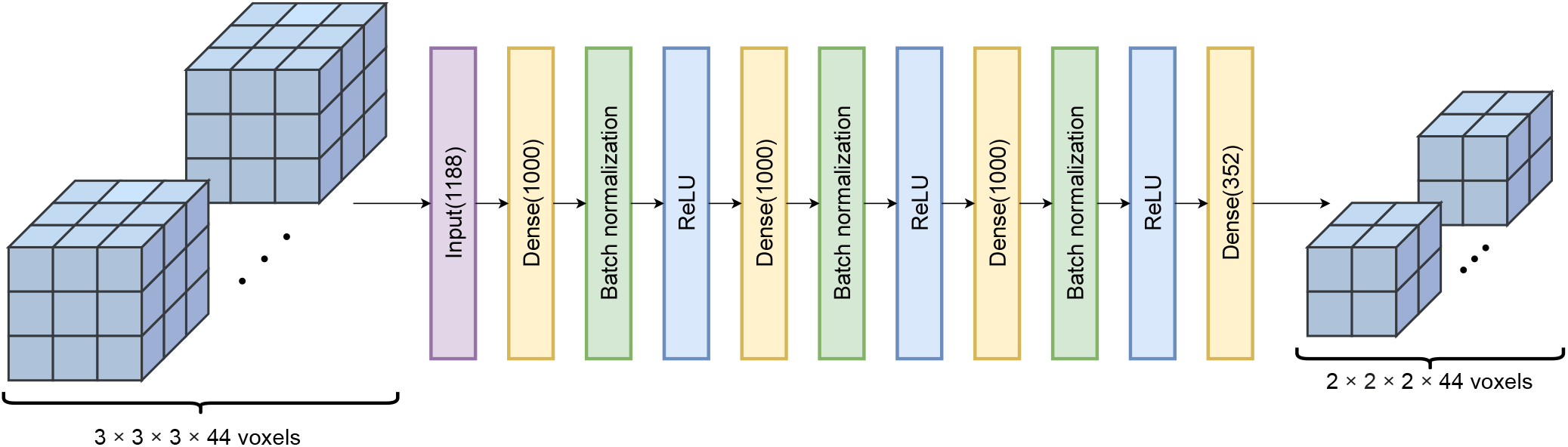
Schematic representation of super-resolution network. The 44 SH coefficients for a 3 × 3 × 3 voxel region of low-resolution data are flattened into a vector and provided as input. The network consists of three blocks, each comprising a dense layer with 1000 nodes, a batch normalization layer, and a ReLU activation. The output is produced by a final dense layer with 352 nodes, which can be rearranged into the 44 SH coefficients of the 2 × 2 × 2 voxel region corresponding to an upsampled version of the central voxel of the input region.

It should be noted that CNNs would be well suited for this upsampling task, due to their efficient use of parameters and the large spatial context that is considered for each predicted output. However, an equivalent CNN implementation to the proposed fully-connected network would require 3-dimensional filters and 44 input channels, which would place very substantial memory requirements on the GPU (unless the CNN works on subvolumes^28^). By comparison, the proposed network has minimal GPU requirements which are fulfilled by most consumer-grade GPUs in the market. Nevertheless, the necessity of providing spatial context in the network input was recognized. We experimented with input regions of size 3 × 3 × 3 and 5 × 5 × 5. However, the larger input region did not provide improved results over the smaller, as the substantial increase in the number of network parameters (from 3.5 to 7.8 million) led to increased overfitting. Henceforth, all results presented are for a network with an input size of 3 × 3 × 3.

### Interpolation

For the purposes of comparison, the low-resolution data were also upsampled using 3D spline interpolation of orders from 0 to 5, where orders 0 and 1 correspond to nearest neighbor interpolation and linear interpolation, respectively. However, it was observed that performance deteriorates with interpolation orders above 2, so results are only shown for orders between 0 and 2, i.e., nearest neighbor, linear, and quadratic. Each spline interpolation operates on one SH coefficient at a time.

The interpolation coordinates were set up in accordance with how the low-resolution data were produced. As each low-resolution voxel is obtained by averaging a 2 × 2 × 2 voxel regions in the high-resolution data, its center is placed at the center of said region. For example, and considering data in a single dimension for simplicity, for high-resolution data with coordinates *x_i_* = *i, i* ∈ {1, *N*}, the corresponding low-resolution data have coordinates *y_j_* = 2*j* – 0.5, *j* ∈ {1,*N*/2}. These latter coordinates are used to set up the interpolation problem to infer the values at the former coordinates. The interpolation was performed using the `ndimage.map_coordinates` function of the `scipy` package for Python.

## Results

### SH metrics

The upsampled high-resolution SHs generated by the proposed diffusion super-resolution network (henceforth referred to as DSR) as well as by spline interpolation were compared to the ground truth high-resolution data on a variety of metrics. In addition to measuring overall performance, it was also measured separately for voxels belonging to WM, GM, and CSF, as specified by the parcellation volume available for each subject.

Figure 2 presents a comparison of mean squared error (MSE, lower is better) and correlation coefficient (higher is better) between the upsampled and ground truth high-resolution data over the SH index, and subdivided by tissue type. The performance for all methods shows a strong dependence on the specific SH coefficient, with four groups of SHs (*Y_ℓm_* with *ℓ*=2, 4, 6, and 8, respectively) that share a roughly similar performance. For MSE, with the exception of the first group of harmonics, performance seems to improve for higher SH coefficients. However, the performance pattern shown by MSE largely reflects the different value ranges that SHs in the four different groups take, with lower values in the ground truth data resulting in similarly low MSE values. In contrast, the correlation metrics, which are independent of the scale of the data, show that performance for all methods diminishes for higher SH coefficients.

**Figure 2.**
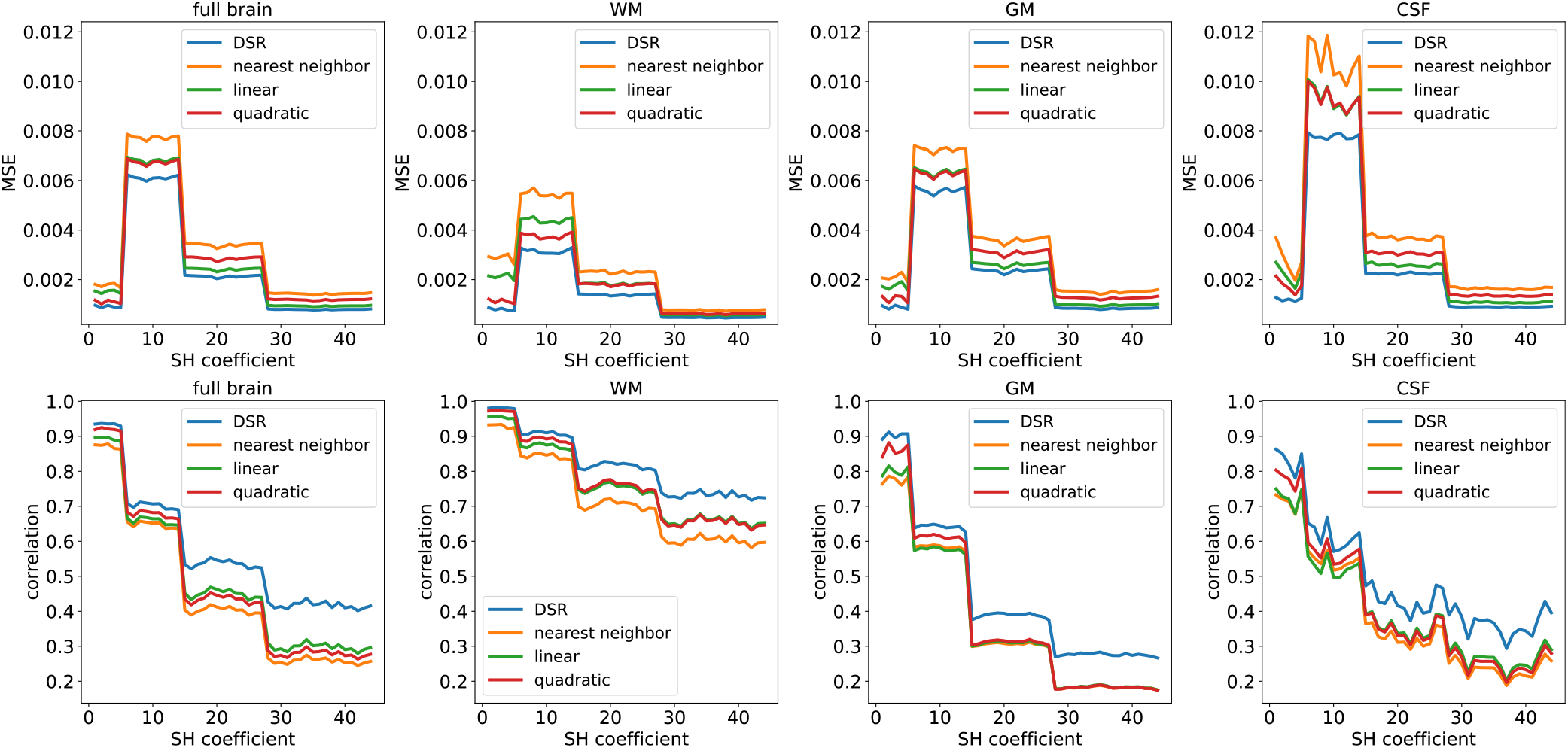
Comparison of similarity metrics between original and upsampled high-resolution SHs, subdivided by metric, tissue type, and SH coefficient. Plotted curves show average metrics for 35 test subjects. Top row: MSE (lower is better). Bottom row: correlation (higher is better). Columns, left to right: overall metrics over the full brain, WM, GM, CSF. MSE: mean squared error, WM: white matter, GM: gray matter, CSF: cerebrospinal fluid.

The performance of all methods also shows a clear dependence on tissue type, with superior performance on WM compared to GM and CSF. This is to be expected, as the highly anisotropic axonal tracts in the WM domain are more self-similar across spatial scales than the shorter dendritic fibers in GM. Furthermore, the free water diffusion seen in CSF does not fulfill the assumptions of diffusion along fiber bundles implicit in CSD, which make the generated fODFs highly random in this tissue type.

Both metrics show that DSR outperforms spline interpolation across all SH coefficients and for all tissue types. For the lowest SH coefficients, corresponding to low angular-frequency components of the fODFs, the gain in performance from using DSR compared to spline interpolation of order 2 is comparable to the gain in performance from increasing the spline order by 1 for splines of order 0 and 1. However, the performance gains are much more substantial for the higher SH coefficients, suggesting that DSR far surpasses the other methods at accounting for the higher angular frequencies of the upsampled fODFs. In addition to MSE and correlation, the various methods were also compared on the basis of mean absolute error, structural similarity index, and peak signal-to-noise ratio, all of which showed DSR outperforming spline interpolation methods (results not shown).

Figure 3 presents difference maps between the ground truth and upsampled (high-resolution) maps of the second SH coefficient (*Y*_2,0_) for a single axial slice of one test subject. As can be seen, the maps produced by DSR show the least difference with respect to the ground truth. Moreover, the difference map from DSR shows the least amount of overt spatial structure, having the appearance of uniform noise, compared to those of the other methods, which suggest that DSR is better at accounting for spatial structures present in the data than the other methods, resulting in sharper upsampled maps.

**Figure 3.**
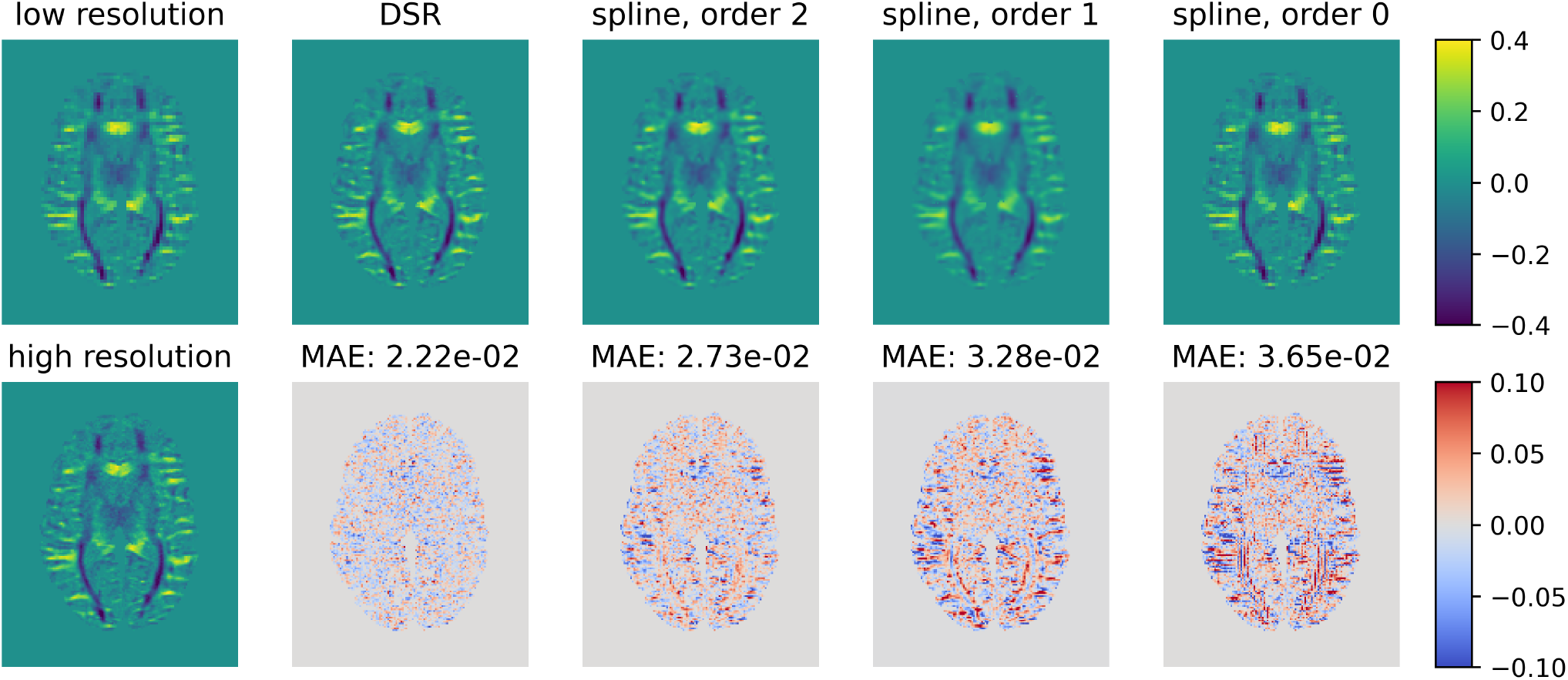
Comparison of upsampling performance of the evaluated methods on the coefficient of the second SH (Y2,0) for a single axial slice of a representative test subject. The leftmost plots show the low-resolution image used as the basis for upsampling and the high-resolution ground truth. Top row: upsampled SH coefficients obtained from the various methods. Bottom row: difference maps between upsampled and ground truth SH maps. MAE: mean absolute error (lower is better). DSR upsampling produces the smallest MAE, as well as having the least amount of overt structure in the resulting difference maps. For the other upsampling methods, certain anatomical structures, such as cortical sulci and gyri, and WM fiber bundles, are recognizable in the difference maps, revealing their uneven performance over different brain features.

### Reconstructed fODF metrics

The upsampled SH coefficients for all methods were converted back into fODFs and compared to the ground truth fODFs. The effects of the different upsampling methods on the estimated local orientations of fiber bundles were studied by extracting the peaks of the fODFs produced by each method, using a relative peak threshold of 0.5, a minimum separation angle between peaks of 25° and specifying a maximum of 5 peaks.

Figure 4 presents several metrics comparing the ground truth and upsampled fODF peaks. Figure 4a shows the average number of fODF peaks in the ground truth and upsampled data for each tissue type. While DSR produces substantially fewer peaks than present in the ground truth for all tissue types, other upsampling methods do so only in WM, and overestimate the number of peaks in GM and CSF. Figure 4b shows the average angle error in the estimated orientation of the main fODF peak for each tissue type. All methods show the best performance on WM, followed by GM and CSF. DSR outperforms spline upsampling methods in all tissue types. Figure 4c shows the fraction of main fODF peaks correctly identified as such by each upsampling method. The performance pattern is similar to previous results, with the best performance seen in WM. Here, too, DSR outperforms spline interpolation methods. Interestingly, the usual performance of the spline methods is reversed for this metric, with splines of orders 0 and 2 giving the best and worst results, respectively.

**Figure 4.**
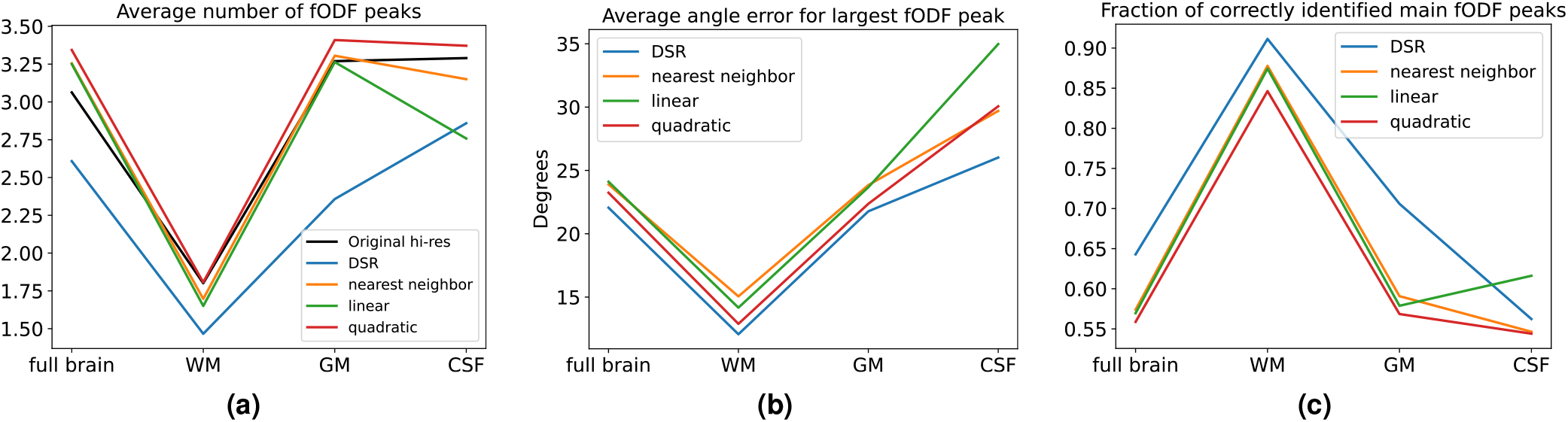
Comparison of fODF peak metrics between original high-resolution data and upsampled data. Plotted curves show average metrics for 35 test subjects. (a) Average number of fODF peaks present in each tissue type. (b) Average angle error in the orientation of the main fODF peak of each fODF. (c) Fraction of voxels where the main fODF peak in the upsampled fODFs is closest in angle to the main peak in the original fODFs.

Figure 5 presents renderings of fODFs for a WM region characterized by crossing fibers. As can be seen, DSR successfully preserves the crossing fibers wherever they are a prominent feature of the data. Furthermore, while there is substantial noise in the ground truth fODFs, with many small peaks present even in areas with a single fiber population, the results of applying DSR are smoother overall, retaining only the most prominent peaks and shrinking the fODFs in CSF regions. Also shown is a rendering of the same region in the low resolution data used as upsampling input, highlighting the increase in gains in spatial specificity of fiber orientation from using DSR.

**Figure 5.**
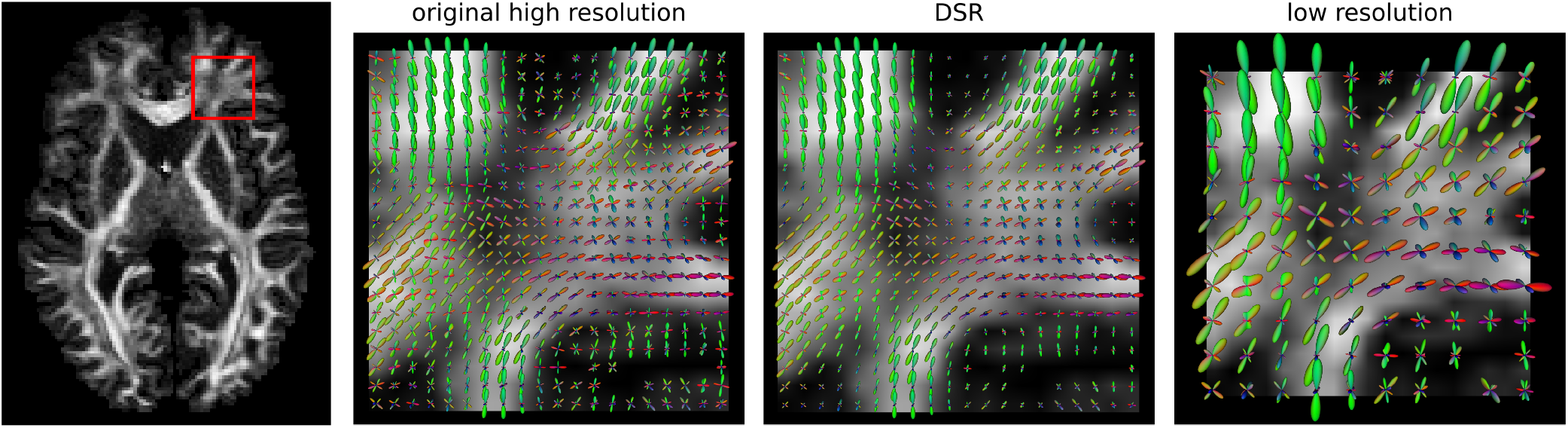
Comparison of fODF glyphs for original high-resolution data and DSR in a region with prominent crossing fibers for a representative test subject. The third plot presents the same region in the low resolution data upsampled by DSR. In addition to correctly representing the main fODF orientations, DSR has a denoising effect on small, spurious fODF components, as well as on regions of uncertain orientation such as CSF.

### Tractography

One of the principal uses of fODFs obtained through CSD involves precise estimation of fiber orientations for tractography, which typically relies on a continuous approximation to the field of fiber orientations. Thus, an accurate estimation of fiber orientations is expected to improve the performance of tractography algorithms. To examine this possibility, we performed deterministic tractography on the original and upsampled high-resolution data for all subjects and evaluated the similarity of the resulting streamlines. Tractography was performed using the EuDX algorithm^35^ in dipy with default parameters and seeds placed on all white-matter voxels. Tracking was stopped when the streamlines exit the white-matter. To facilitate comparison between streamlines, only the direction of the principal peak was traced in voxels with multiple fODF peaks.

For each seed point, we calculated the Euclidean distance between the streamline endpoints obtained from the original data and data upsampled with each of the tested methods. This resulted in mean distances over subjects and seedpoints of 15.9 for

DSR, 16.8 for quadratic splines, 17.6 for linear interpolation, and 17.7 for nearest neighbor interpolation. The improvement achieved by DSR over quadratic spline interpolation is comparable to the improvement the latter produces over nearest neighbor interpolation. These results suggest that upsampling fODF data with DSR prior to performing tractography may indeed result in more accurate streamline tracking.

### Repeated upsampling

In order to explore the limitations of DSR upsampling, as well as to evaluate its performance at upsampling resolutions it was not specifically trained on, we experimented with upsampling the original high-resolution data with DSR. To this end, the data was upsampled repeatedly, by feeding the output of DSR back as input. This upsampling was repeated three times, being very computationally-demanding. As there is no ground truth for the super-resolved data, we can only perform a qualitative evaluation of the results. Figure 6 presents plots of the magnitudes of the first three fODF peaks, as well as of the overall number of peaks per voxel, for the super-resolved data. Repeated upsampling does not result in blurring of the data, as could be expected, but actually produces sharp spatial maps with a high level of detail, although showing some streaking and checkerboard artifacts on the higher resolutions. Some notable features can be seen in Figure 6d, where long, single-voxel-wide regions of crossing fibers are well-resolved under repeated upsampling, with some features retaining a width of a single voxel in the highest achieved resolution.

**Figure 6.**
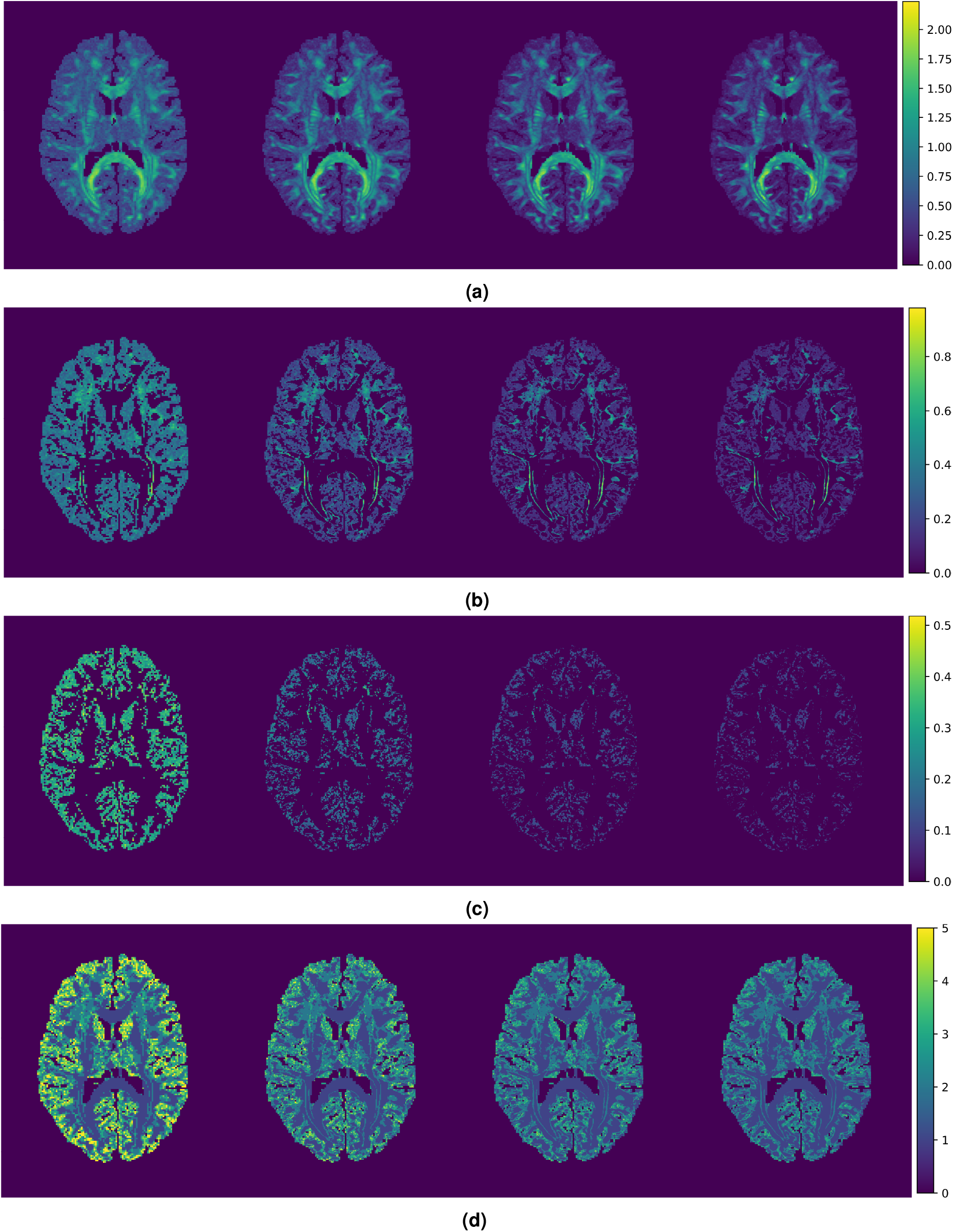
Results of repeated upsampling with DSR of original high-resolution data. The leftmost volume in each row is produced from the original high-resolution data, with each subsequent volume having double the resolution in each axis of the previous. (a) Magnitude of first fODF peak. (b) Magnitude of second fODF peak, where present. (c) Magnitude of third fODF peak, where present. (d) number of fODF peaks per voxel. In all cases, DSR is capable of generating sharp results with a remarkable level of anatomical detail. Voxel resolutions, from left to right: 1.25 mm, 0.625 mm, 0.313 mm, 0.156 mm.

The denoising effects of DSR are also made apparent under repeated upsampling. It can be seen that the magnitude of the first and second fODF peaks is generally preserved or somewhat amplified in WM, where it is well-specified. On the other hand, the magnitudes of all fODF peaks are attenuated with each upsampling in regions outside WM, where the assumptions of CSD do not fully apply, and the self-similarity of diffusion orientation across scales is lower. This is especially apparent for the third fODF peak, which is rarely detected in WM, and is almost completely suppressed after upsampling three times. The overall number of fODF peaks per voxel is also reduced, as shown in Figure 6. It should also be noted that a reduction in the number of fODF peaks is to be expected as the resolution of data increases, since large voxels showing multiple crossing fibers can increasingly be resolved into their component fibers.

## Discussion

Throughout the presented experiments, DSR has shown improved upsampling performance over spline interpolation methods (see Figures 2, 3, 4a, 4c). However, the learning-based nature of DSR’s upsampling, coupled with the large training corpus used, gives it substantial advantages over ordinary interpolation methods that go beyond the raw metrics. First, it allows DSR to conform its upsampling approach to local anatomical brain features, resulting in sharper upsampled maps (see Figure 3). Furthermore, it allows DSR to distinguish those aspects of the data which are coherent across its training dataset, such as the orientations of the main fODF peaks, from those that are not coherent, such as spurious fODF peaks caused by the high noise-susceptibility of CSD. This latter feature causes DSR to have a denoising effect on the upsampled fODFs, allowing it to remove spurious fODF components that are present in the high-resolution ground truth while preserving the relevant orientation information (see Figure 5). Remarkably, it is capable of doing this while using only 1/8 of the data available in the ground truth. A possible explanation for this is that our DSR approach uses the information in all 44 SH coefficients concurrently, while the spline interpolation operates on one SH coefficient at a time.

These advantages are especially apparent when DSR is tested on repeated upsampling, where it has been shown to produce sharp scalar maps with a remarkable level of spatial detail (see Figure 6). Although the extent to which these results correspond to real brain anatomy is uncertain due to the lack of ground truth, it is nevertheless notable that the performance of DSR does not show a substantial diminishment even after three iterative applications, which results in a voxel size of 156 μm. Here, our underlying assumption is that the transformation that maps the fODFs at a lower-resolution of the downsampled data to the resolution of the acquired data is the same as the transformation that would be employed to obtain smaller and smaller voxel size. Such *self-similarity* persists across all length scales for fractal media. Indeed, fractal-concepts have been employed in profiles of neural cells obtained via microscopic methods,^36^ the contour of the brain cortex measured via MRI,^37^ and finally for modeling a power-law scaling encountered in the measurements of magnetic resonance measurements of time-dependent diffusion.^38, 39^

We note that our approach can be extended and generalized in a number of ways. The training performed on the patches of fODFs could be performed on diffusion-weighted data,^27^ which could feature other sampling schemes such as those that cover a larger region of the sampling space.^13^ A basis other than the SH basis could be employed for representing such data.^40^ The input and/or the output could include scalar measures of tissue structure.^41^ A representation of asymmetric ODFs^42–44^ could be employed that could allow disambiguation of asymmetric features that are encountered for bending fibers or Y-shaped crossings. Regarding using a larger spatial region of interest (5 x 5 x 5), this can possibly be achieved by adding Dropout regularization between the dense layers to prevent overfitting.

## Conclusion

We have presented a deep-learning-based super-resolution approach for upsampling diffusion fODF data. The proposed method outperforms standard spline interpolation methods, and is capable of rendering nuanced anatomical features in scalar maps derived from the fODF data. We believe the proposed method can be particularly useful for improving tractography, as it inherently relies in interpolation for estimating the local orientation of fiber tracts in positions between the available voxels.

## Acknowledgements

We thank Gozde Unal and Enes Albay for stimulating discussions. This research was supported by the ITEA/VINNOVA project ASSIST (Automation, Surgery Support and Intuitive 3D visualization to optimize workflow in IGT SysTems, 2021-01954).

## Author contributions statement

D.A. implemented the neural networks and performed all the analyses. E.Ö. conceptualized the study. A.E. and E.Ö. supervised the work. All authors wrote the manuscript.

